# Simultaneous ribosome profiling of human host cells infected with *Toxoplasma gondii*

**DOI:** 10.1101/619916

**Authors:** Michael J. Holmes, Premal Shah, Ronald C. Wek, William J. Sullivan

## Abstract

*Toxoplasma gondii* is a ubiquitous obligate intracellular parasite that infects the nucleated cells of warm-blooded animals. From within the parasitophorous vacuole in which they reside, *Toxoplasma* tachyzoites secrete an arsenal of effector proteins that can reprogram host gene expression to facilitate parasite survival and replication. Gaining a better understanding of how host gene expression is altered upon infection is central for understanding parasite strategies for host invasion and for developing new parasite therapies. Here, we applied ribosome profiling coupled with mRNA measurements to concurrently study gene expression in the parasite and in host human foreskin fibroblasts. By examining the parasite transcriptome and translatome, we identified potential upstream open reading frames that may permit the stress-induced preferential translation of parasite mRNAs. We also determined that tachyzoites reduce host death-associated pathways and increase survival, proliferation, and motility in both quiescent and proliferative host cell models of infection. Additionally, proliferative cells alter their gene expression in ways consistent with massive transcriptional rewiring while quiescent cells were best characterized by re-entry into the cell cycle. We also identified a translational control regimen consistent with mTOR activation in quiescent cells, and to a lesser degree in proliferative cells. This study illustrates the utility of the method for dissection of gene expression programs simultaneously in parasite and host.

**Importance:** *Toxoplasma gondii* is a single-celled parasite that has infected up to one-third of the world’s population. Significant overhauls in gene expression in both the parasite and the host cell accompany parasite invasion, and a better understanding of these changes may lead to the development of new therapeutic agents. In this study, we employed ribosome profiling to determine the changes that occur at the levels of transcription and translation in both the parasite and the infected host cell at the same time. We discovered features of *Toxoplasma* mRNAs that suggest a means for controlling parasite gene expression under stressful conditions. We also show that differences in host gene expression occur depending on whether they are confluent or not. Our findings demonstrate the feasibility of using ribosomal profiling to interrogate the host-parasite dynamic under a variety of conditions.

## Introduction

*Toxoplasma gondii* is a ubiquitous intracellular parasite that infects nucleated cells of warm-blooded animals. Upon infection, tachyzoites undergo a period of rapid replication and dissemination throughout the host (1). Usually without producing symptoms, host immunity induces conversion of tachyzoites into latent bradyzoites that are contained within tissue cysts that can persist for the life of the host (2). During immunosuppression, bradyzoites can reconvert into tachyzoites, causing severe disease due to lysis of host cells in critical organs and tissues (3). There is currently no treatment available to eradicate tissue cysts, and the frontline anti-folate drugs that control acute infection exhibit serious toxic effects in patients (4). A better understanding of how intracellular tachyzoites interact with their host cells may reveal new drug targets to treat this pathogen.

It is now well established that tachyzoites induce significant changes inside their host cell. An arsenal of parasite proteins is released via secretory organelles at the apical end of the parasite upon invasion as the parasites establish the non-fusogenic parasitophorous vacuole in which they reside and replicate (5). For example, secreted proteins ROP16, GRA6, GRA16, and GRA24 modulate transcription factor activity in the host cell, inducing gene expression that is thought to favor parasite proliferation (reviewed in (6)).

In addition to initiating transcriptional changes, tachyzoites are suggested to modulate mRNA translation in the infected host cell. *Toxoplasma*-infected host cells show increased molecular target of rapamycin (mTOR) activity and subsequent phosphorylation of downstream mTOR targets (7–9). Tachyzoite-induced mTOR activation is suggested to increase translation of selected gene transcripts (9). *Toxoplasma* infection may also activate the eukaryotic initiation factor-2α (eIF2α) kinase GCN2 in MEF cells in response to tachyzoite consumption of host cell arginine (10). Phosphorylation of eIF2α lowers translation initiation, which serves to conserve nutrients and energy, and enhances the translation of select mRNAs involved in stress adaptation (11). *Toxoplasma* also possesses a complement of protein kinases that phosphorylate the parasite eIF2α in response to stress and inducers of bradyzoite development (reviewed in (3)). Collectively, these studies suggest that translational control mechanisms are operating in both the parasites as well as in their host cells during infection.

Ribosome profiling (Ribo-seq) has emerged as a powerful method for analyzing translational dynamics quantitatively and qualitatively (12). This technique has been applied to protozoan parasites to improve genome annotation and to investigate translational control in different parasite life stages or in response to chemotherapeutics (13–18). In the cases of *Plasmodium falciparum* (the parasite that causes malaria) as well as *Toxoplasma gondii* intracellular and extracellular tachyzoites, these experiments helped to identify translational control mechanisms suggested to involve upstream open reading frames (uORFs) (13, 14).

In this study, we present a method that can be used to interrogate translational control and transcriptome changes in intracellular parasites and their host cells simultaneously. Using Ribo-seq, we show that one can profile intracellular tachyzoite and host cell transcriptomes and translatomes concurrently. We utilize proliferative and quiescent host cell models of infection to determine growth-independent changes that occur in response to infection. We determine that gene expression changes in proliferating host cells are largely driven by changes of mRNA levels. By comparison, quiescent host cells display changes in gene expression that are controlled at the levels of the transcriptome and translatome that are consistent with re-entry into the host cell cycle. However, bulk translational capacity, defined as the relative amount of translating ribosomes, remains largely unaltered in the infected host cell under both growth conditions. We also leveraged our datasets to investigate the tachyzoite translatome for potential uORFs, which can be regulators of transcript-specific translational control. Our findings suggest that concurrent Ribo-seq will be a valuable tool to identify changes in both host cells and parasites under a wide-variety of experimental conditions.

## Results and Discussion

### Concurrent host-parasite ribosome profiling

Confluent HFF monolayers are used routinely to study *Toxoplasma* infection and host-parasite interactions *in vitro* whereas subconfluent cells are frequently used in studies of stress-induced translational control in other systems. Since primary fibroblasts become quiescent upon reaching confluency and subconfluent fibroblasts are proliferative (19), we reasoned that each growth condition would be a useful model for examining the growth-dependent and -independent gene expression changes that occur at the transcriptome and translatome levels in host cells upon tachyzoite infection. We assessed bulk translation in confluent and subconfluent HFFs by polysome profiling, which gauges translation by comparing the relative contribution of polysomes and monosomes to the profile (Fig. 1A-B). Confluent HFFs have low basal translation, as evidenced by a prominent monosome peak and modest polysomes (Fig. 1A, solid lines). By comparison, proliferating subconfluent HFFs display larger polysomes relative to the monosome peak, which is indicative of more robust translation (Fig. 1A, dotted lines). We quantified the proportion of RNA involved in the polysome fraction as an indication of overall bulk translation (Fig. 1B). Indeed, translation is significantly more robust in subconfluent compared to confluent HFFs. There were modest, albeit statistically significant, changes in the polysome profiles following 24 hours of infection with tachyzoites (Fig. 1B). These changes may be a reflection of altered translational control in the host cell and/or represent a contribution of parasite RNA to the profiles.

**Figure 1.**
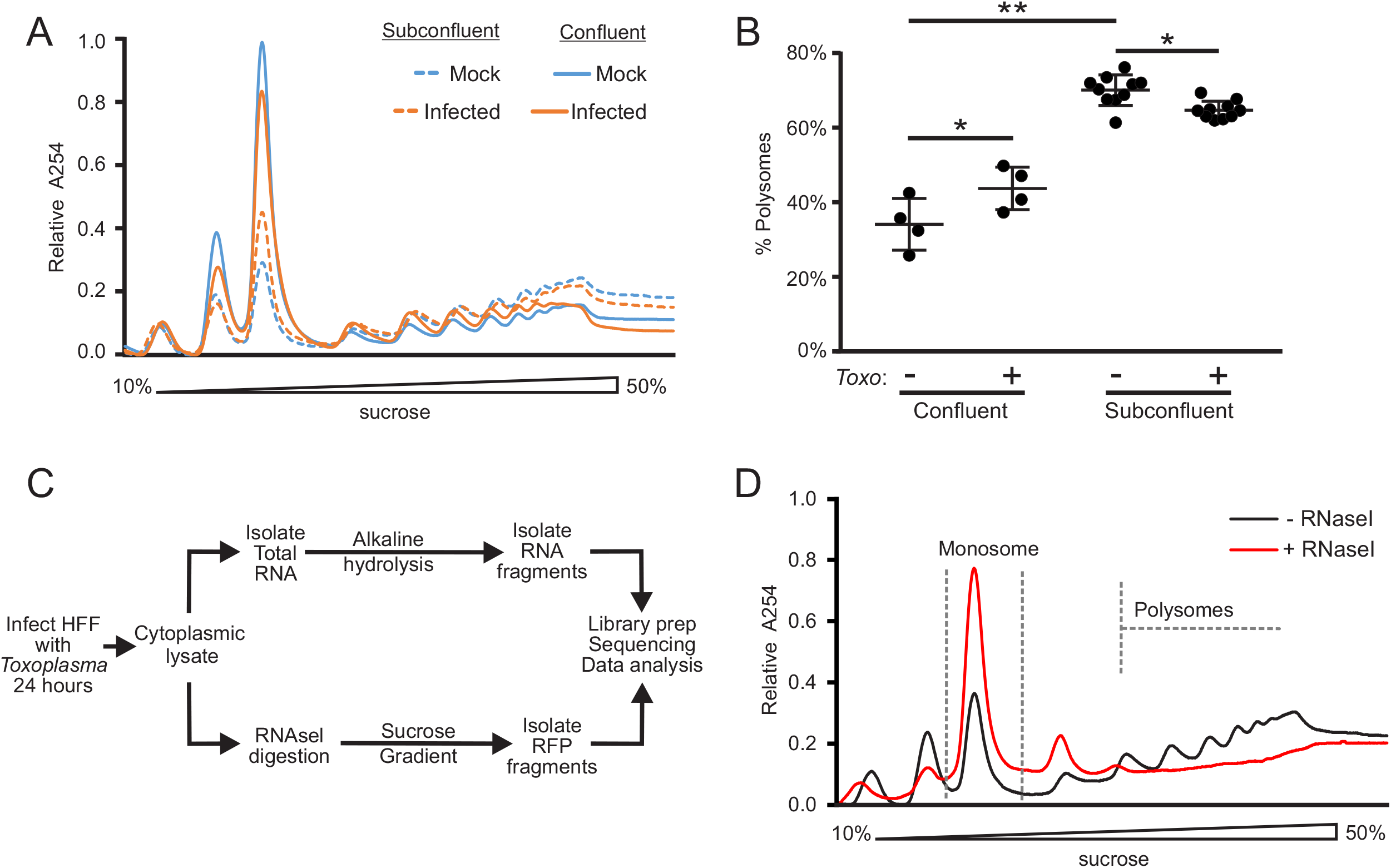
Polysome and ribosome profiling of *Toxoplasma*-infected HFFs. A) Representative examples of polysome profiles from mock-infected or tachyzoite-infected confluent (solid lines) and subconfluent (dotted lines) HFFs 24 hours after infection with RH strain tachyzoites (MOI~10). B) Polysome abundance analysis of profiles performed as in (A). Median proportions of polysomes and standard deviation are presented for each condition. C) Experimental design of ribosome profiling experiments. HFFs were infected as in (A) and cytoplasmic lysate was partitioned for subsequent RNA-seq and ribosome profiling analysis. Total RNA was fragmented by alkaline hydrolysis. For ribosome profiling, cytoplasmic lysate was digested with RNaseI, and monosomes were collected after centrifugation on sucrose gradients. RFPs were collected and processed along with small RNA fragments for sequencing library preparation. D) Example of cytoplasmic lysate generated from infected subconfluent HFFs with and without digestion by RNaseI. Grey dashes outline the monosome fraction of the gradient collected for RFP isolation. *P ≤ 0.05; **P ≤ 0.001, one way ANOVA with Tukey comparison.

We conducted our ribosome profiling experiments 24 hours after infection with RH strain tachyzoites using a protocol modeled after previously published techniques (12, 13). A schematic diagram outlining the experimental design is shown in Figure 1C. The infected HFF cells were treated with cycloheximide (CHX) to stop translational elongation and then lysed in cytoplasmic lysis buffer. An aliquot of the cytoplasmic extract was collected for RNA-seq analysis; the remainder was digested with RNase I followed by sucrose density centrifugation. The RNase I digests mRNA regions that are not occupied by ribosomes, allowing for only the ribosome protected fragments, or footprints (RFPs), to be collected in the resulting monosome peak after sucrose density centrifugation (Fig. 1D). The peak corresponding to the monosome was collected (Fig. 1D) and the resulting RFPs were used for subsequent sequencing library preparation (Fig. 1C). In our experiments, sucrose density centrifugation as described (13) reduced downstream rRNA contamination in our sequencing libraries compared to the sucrose cushion method of RFP collection previously described (data not shown) (12).

We mapped our sequencing data independently to the human and *Toxoplasma* ME49 strain transcriptomes. Although our experiment was conducted with the type I strain RH, all of our analyses were mapped to the type II reference strain ME49 because it is the most thoroughly annotated of all strains available in the ToxoDB. Metagene profiles were generated for each library and the profile analyses revealed that a large portion of the RFPs mapped to the annotated coding sequence (CDS) start codons and progressively decreased immediately downstream in both infected human host cells and intracellular tachyzoites (Fig. 2A-B). This “ramp-like” pattern is reminiscent of that observed in studies detailing the effects that CHX contributes to RFP distribution patterns when used in Ribo-seq experiments (20–22). Our results are consistent with the idea that CHX causes the same “ramp-like” perturbation to *Toxoplasma* ribosome profiles as reported in yeast and mammalian cells and can be leveraged to facilitate the identification of translated uORFs in tachyzoites since it accentuates RFP peaks near start codons.

**Figure 2.**
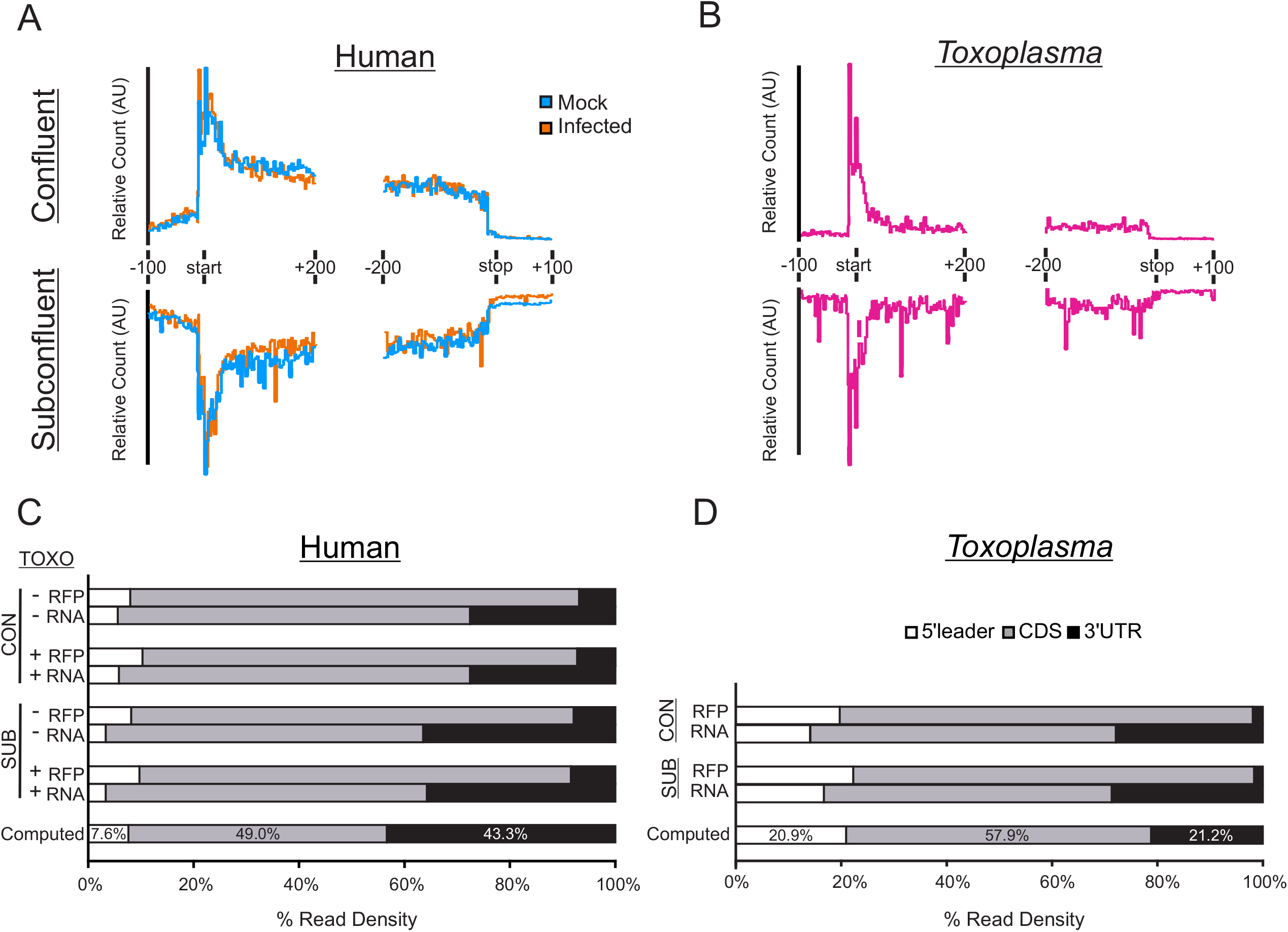
Enrichment of RFPs on human and *Toxoplasma* coding sequences. A) Metagene plots of ribosome profiling data from mock-infected and tachyzoite-infected HFFs. Transcript diagrams are labelled and drawn proportionally. Plots are generated from a representative replicate. B) Representative metagene plot as in (A) for *Toxoplasma*-mapping data. C) Quantitation of ribosome profiling (RFP) and RNA-seq (RNA) reads that map to each transcript feature. A computed model transcript, representing the average contribution of each feature to transcript length (expressed as a percentage of total transcript) is shown for comparison. D) Quantitation of RFP and RNA reads as in (C) for the sequencing reads that mapped to *Toxoplasma* sequences. CON: data from confluent host cell experiment; SUB: data from subconfluent host cell experiment.

The metagene profiles also revealed a consistently higher ribosome density along the 5’-leader sequences in both host and parasite cells compared to the 3’-UTRs of the transcripts (Fig. 2A-B). In analyses of the *P. falciparum* translatome during the intraerythocytic developmental cycle, an enrichment for 5’-leader sequence representation was noted and attributed to ribosomal occupancy but did not correlate with the presence or location of uORFs (13, 23). To determine if a similar phenomenon occurs in *Toxoplasma*, we first computed the average contribution of each annotated feature (i.e., 5’-leader, CDS, and 3’-UTR) to coding mRNAs transcriptome-wide using data from the *Toxoplasma* and human genome annotations EuPathDB (24) and data generated in (25). With this information, we constructed a model for the average computed transcript in host and parasite (Fig. 2C-D). The average human 5’-leader composes 7.6% of the total transcript length whereas this feature contributes 20.9% of the total *Toxoplasma* mRNA length. We then quantified the ribosome density along all coding transcripts in both host cells and parasites, segregating them according to their feature annotation (Fig. 2C-D). Results indicate that 8-10% of human RFPs and 20-22% of *Toxoplasma* RFPs mapped to 5’-leaders, suggesting that the rate of RFP mapping to 5’-leaders in both organisms is proportional to their length. Since the rate of RFP mapping to 5’-leaders in both organisms appeared proportional to their length, and the accumulation of RFPs within 5-’leaders has also been attributed to the CHX treatment (22), we suggest that we are also observing CHX-induced ribosome occupancy along the 5’-leaders. Moreover, this result appears to be enhanced in *Toxoplasma* compared to human cells due to the longer 5’-leaders in the parasite. In contrast, we saw enrichment of RFPs mapping to CDS regions and a decreased representation of 3’-UTRs in all conditions. Mapping of the RNA-seq data did not show a similar enrichment (Fig. 2C-D). Approximately 80% of human and ~75% of *Toxoplasma* RFP reads mapped to CDS regions and ~7-8% of human and ~2% of *Toxoplasma* RFPs mapped to 3’-UTRs. The enrichment of RFPs in the CDS is expected of Ribo-seq experiments, indicating that we have substantial data to assess translatome changes and potential translational control in both infected host cell and in the parasite.

### Ribo-seq analysis of intracellular tachyzoites in HFF host cells

In our infected host HFFs, ~15-25% of the sequences mapped to *Toxoplasma* (Table 1), making it possible to detect parasite-specific sequences for a survey of the tachyzoite translatome. We assessed the ribosome density, a measure of translation, for all coding transcripts that met our filtering criteria, which consisted of coding genes that have annotated 5’- and 3’-UTRs and a normalized read count ≥ 5 in each of our biological triplicates as determined by DESeq2 (26). For tachyzoites grown in confluent HFFs, this analysis resulted in 4,453 transcripts, whereas 5,762 transcripts met our cutoff for tachyzoites cultured in subconfluent HFFs (Fig. 3A, Table S1A-B). Bulk ribosomal density was calculated for each transcript by taking the quotient of normalized RFP and RNA-seq reads. There was no statistically significant difference between the two ribosomal density profiles (Student’s *t*-test, P = 0.07), indicating that the tachyzoite translational machinery functions at a similar capacity and throughput regardless of host cell confluency (Fig. 3A).

**Figure 3.**
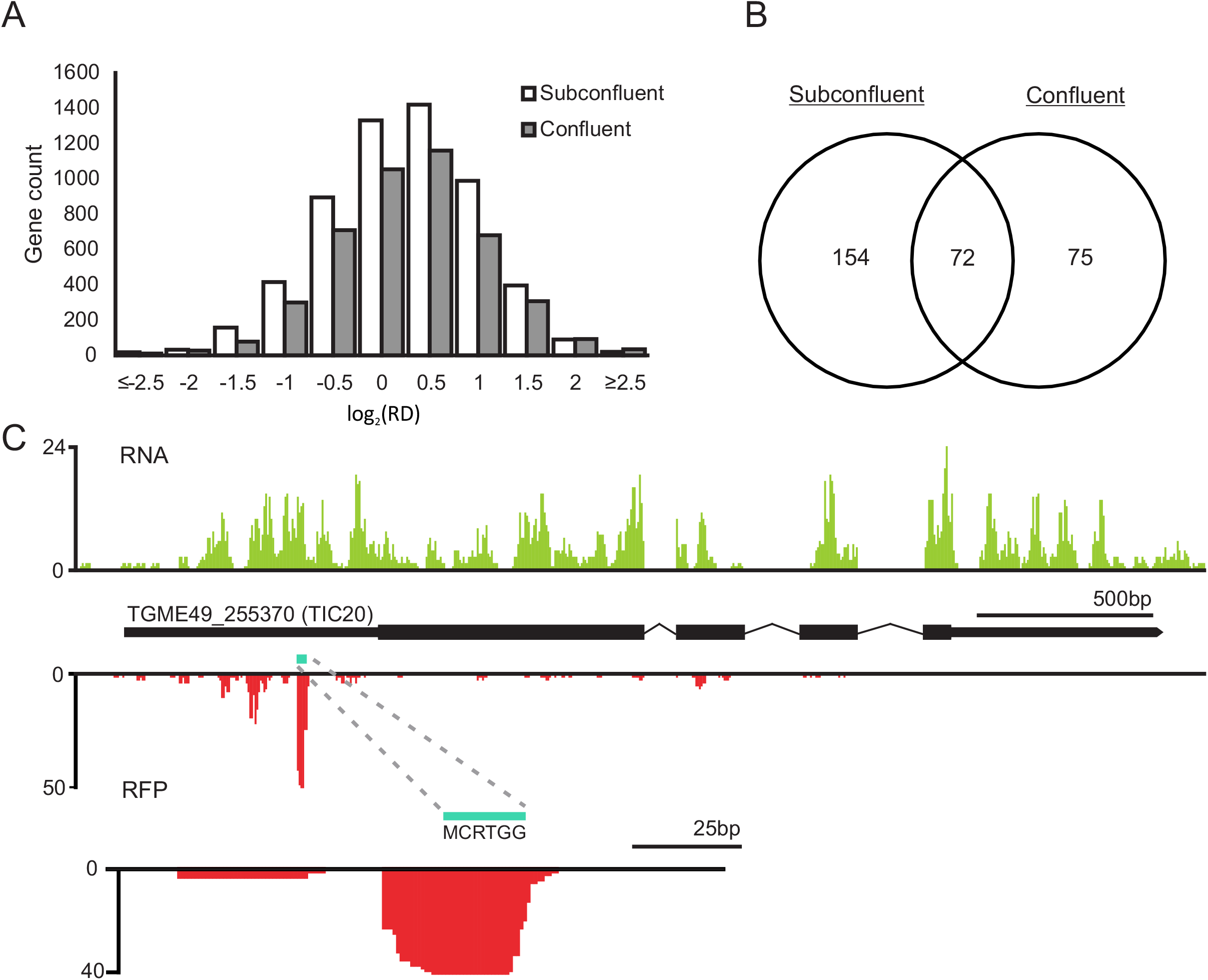
The *Toxoplasma* tachyzoite translatome and potential uORFs. A) Distribution of tachyzoite ribosome density (RD) transcriptome-wide between parasites grown in confluent or subconfluent HFFs for 24 hours. B) Venn diagram of transcripts identified as encoding potential uORFs from tachyzoites grown in confluent or subconfluent HFFs. Transcripts with at least 2-fold greater ribosome density in their annotated 5’-leader than their annotated CDS were included in the analysis. C) Potential uORF in the 5’-leader of TIC20. RNA-seq data is displayed above the gene model. Ribosome profiling data are positioned below the gene model, showing a peak centered over a potential uORF that encodes a six amino acid peptide.

**Table 1.** Ribosome profiling and RNA-seq library mapping statistics.

| Sample | Confluent |  |  |  | Subconfluent |  |  |  |
| --- | --- | --- | --- | --- | --- | --- | --- | --- |
|  | Human |  | *Toxoplasma* |  | Human |  | *Toxoplasma* |  |
|  | RFP | RNA | RFP | RNA | RFP | RNA | RFP | RNA |
| HFF1 | 63.49% | 86.65% | - | - | 63.03% | 80.64% | - | - |
| HFF2 | 53.38% | 87.89% | - | - | 63.89% | 82.35% | - | - |
| HFF3 | 68.22% | 86.86% | - | - | 60.44% | 80.76% | - | - |
| HFF-RH1 | 60.74% | 64.68% | 25.23% | 16.58% | 50.46% | 56.23% | 14.67% | 21.97% |
| HFF-RH2 | 43.47% | 66.97% | 18.30% | 17.71% | 48.79% | 55.16% | 21.51% | 23.95% |
| HFF-RH3 | 68.33% | 68.85% | 14.15% | 14.54% | 45.39% | 55.25% | 19.99% | 22.70% |

Several reports have described the mechanisms of translational control in *Toxoplasma*, especially through the involvement of the parasite integrated stress response (ISR) pathway (reviewed in (3)). For example, our lab identified a subset of transcripts that were preferentially translated in tachyzoites upon induction of ER stress, but the mechanism of preferential translation was not elucidated (27). One potential mechanism could be through the usage of functional uORFs in *Toxoplasma* since these *cis*-acting elements provide a means to achieve the transcript-specific upregulation of stress-responsive proteins during ISR activation hypothesized to be necessary for bradyzoite formation (3, 28). The *in silico* prediction of uORFs is complicated by the unusually lengthy 5’-leaders of *Toxoplasma* transcripts (14, 29) (Fig. 2D). Consequently, longer 5’-leaders are inherently more likely to contain ORFs upstream of the annotated CDS (14).

Ribosome profiling facilitates uORF discovery since it provides evidence of ribosome occupancy along distinct ORFs. In their Ribo-seq experiment of intracellular and extracellular tachyzoites, Hassan *et al*. reported evidence of 2,770 uORFs across the transcriptome that generally displayed low ribosomal coverage (14). We also leveraged our dataset to discover potential uORFs in our intracellular tachyzoite samples. We identified all coding transcripts from our dataset with at least a two-fold higher ribosome density on their annotated 5’-leaders than their CDS, reasoning that uORFs involved in stress-induced translational control are often translated at the expense of their downstream CDS (28). This analysis yielded 226 and 147 transcripts for tachyzoites grown in subconfluent and confluent HFFs, respectively, with 72 transcripts present in both culture conditions (Fig. 3B; Table S2A-B). Importantly, our Ribo-seq experiments were conducted using a mixture of HFF and intracellular tachyzoites instead of purified parasites, and our confluent HFF experiment had a lower parasite contribution than our subconfluent experiment (Table 1). Consequently, the depth of coverage for our studies was less than that reported in Hassan *et al*. (14), likely explaining the lower number of hits in our dataset.

We visually inspected the ribosome profiles of the transcripts with high 5’-leader ribosomal occupancy for regional spikes in RFP coverage that overlapped putative uORFs. A potential uORF was identified in the transcript for TIC20, a member of the apicoplast protein translocation machinery (30). The TIC20 gene was well-mapped over the entire transcript by RNA-seq, yet displayed a strong RFP peak in its 5’-leader and very low levels of CDS translation (Fig. 3C). A small uORF encoding a hexapeptide is overlapped by Ribo-seq reads that display a characteristic 12 nucleotide offset from the start codon typical of ribosome footprints (12), suggesting this to be a functional uORF. Concurrent ribosome profiling experiments can be designed to validate and/or screen for potential uORFs in stressed intracellular parasites in the future. Determination and validation of functional uORFs will likely reveal cases of transcript-specific translational control that are involved in adaptation of *Toxoplasma* to stressful stimuli, including bradyzoite induction conditions.

### Proliferating and quiescent host cells respond differently to tachyzoite infection

Upon invasion, *Toxoplasma* tachyzoites remodel the host cell environment by recruiting host organelles such as the ER, Golgi apparatus, endo-lysosomes, and mitochondria to the parasitophorus vacuole (31–33). Along with the physical reorganization of the infected host cell are concomitant changes in host gene expression, some of which are mediated directly by secreted parasite effectors originating from rhoptries and dense granules proteins (reviewed in (6)). Research has also revealed that different types of host cell backgrounds respond to *Toxoplasma* in different ways (34). However, the degree to which proliferating and quiescent host cells vary in their response to infection remains to be investigated. Therefore, we profiled the transcriptomes and translatomes of confluent and subconfluent HFFs that had been infected (or mock-infected) with *Toxoplasma* for 24 hours. We identified 8,461 genes that met our inclusion criteria (detected at both the RFP and RNA levels with a normalized mean count ≥ 5) in confluent fibroblasts, of which 1,016 were differentially expressed with a log_2_ fold change (FC) ≥ ±1 and met a false discovery rate (FDR) of 0.01 at either the RNA and/or RFP levels (Table S3A). By the same metrics, we identified 904 differentially expressed genes among the 10,164 detectable genes in subconfluent cells (Table S3B). A total of 8,427 genes were detected consistently in all samples (Fig. 4A). We noted most parasite-induced dysregulated expression appeared to be equally changed at the transcriptome and translatome levels in both confluent and subconfluent cells. This finding suggests that these genes constitute a core, shared transcriptional response to tachyzoite infection. Gene ontology analysis (35) of the downregulated genes suggests an enrichment (FDR = 0.05) of membrane-resident G-protein coupled receptors in subconfluent cells whereas, upregulated genes were enriched for cytokines and transcriptional machinery (Table 2). Confluent HFFs also displayed a high enrichment suggestive of transcription factors and associated machinery (Table 3). Highlighting the fidelity of our dataset, we saw changes in host gene expression that have been reported previously, such as the parasite-induced upregulation of CCL11, HAS3, ATF3, and EGLN3, all genes that were included as host response factors on the ToxoGeneChip (36).

**Figure 4.**
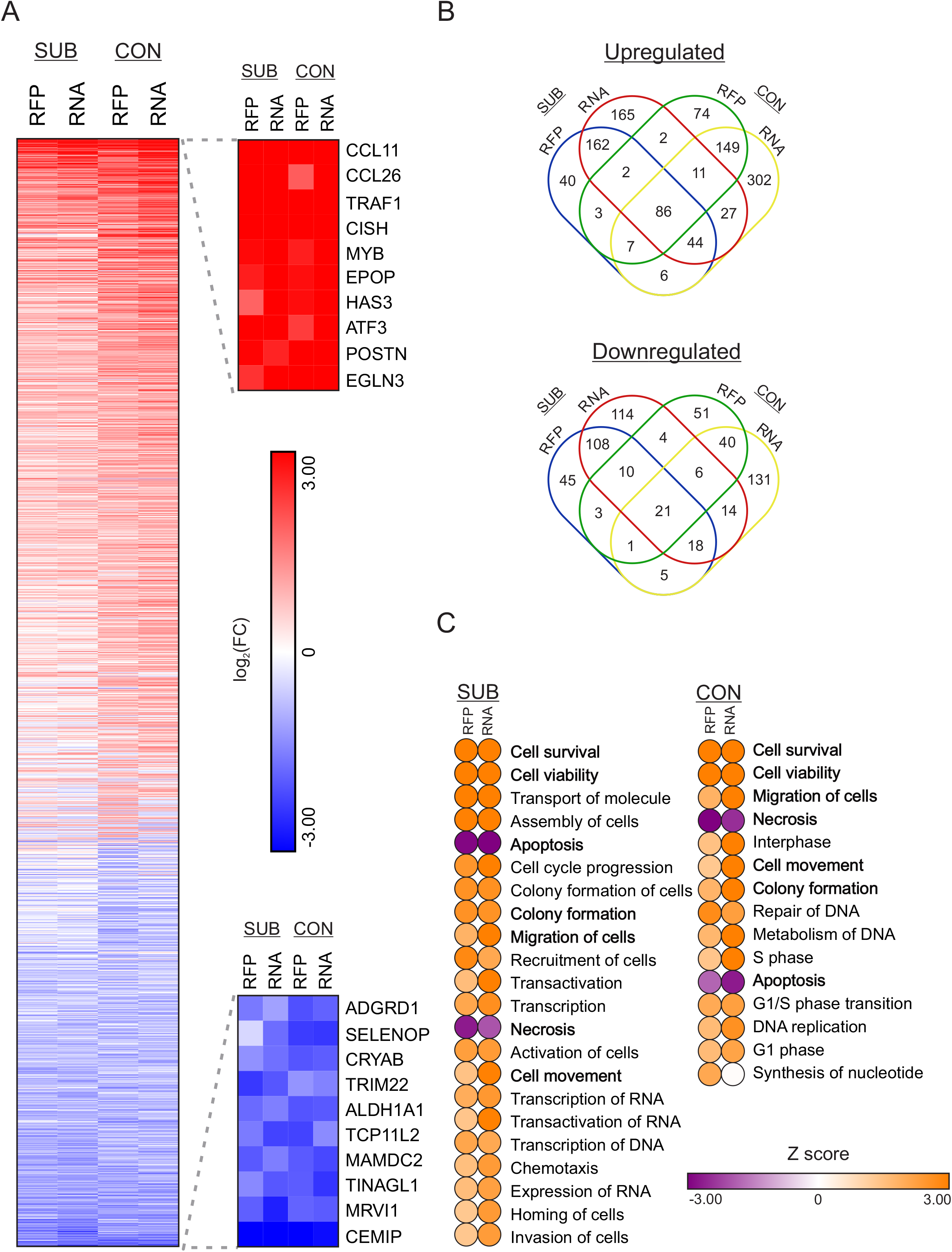
Differentially regulated genes in HFF host cells following 24 hours of tachyzoite infection. A) Heat map of all genes identified in ribosome profiling (RFP) and RNA-seq (RNA) datasets. Lanes are segregated by subconfluent (SUB) and confluent (CON) sequencing libraries. Data represents log_2_ transformed fold change upon infection. Rows are ordered from most induced (red) to most repressed (blue). The top ten genes from each extreme are presented. B) Venn diagrams showing the relationship between shared and growth condition-specific up- and downregulated genes upon infection. C) Ingenuity Pathway Analysis of modulated cellular and molecular functions in subconfluent and confluent HFFs upon tachyzoite infection. Pathways with a significance of P ≤ 0.05 and displayed a Z score ± 2 in either the RFP or RNA datasets are shown. Pathways that are common between confluent and subconfluent HFFs are in bold.

**Table 2.** Molecular function enrichment of gene expression changes in subconfluent HFFs upon tachyzoite infection.

| Category | Term | Padj |
| --- | --- | --- |
| Upregulated |  |  |
| GO:0005125 | cytokine activity | 6.60E-04 |
| GO:0005539 | glycosaminoglycan binding | 1.98E-03 |
| GO:0008201 | heparin binding | 4.55E-03 |
| GO:1901681 | sulfur compound binding | 6.33E-03 |
| GO:0046873 | metal ion transmembrane transporter activity | 1.14E-02 |
| GO:0000981 | RNA polymerase II transcription factor activity, sequence-specific DNA binding | 1.90E-02 |
| GO:0004866 | endopeptidase inhibitor activity | 4.61E-02 |
| Downregulated |  |  |
| GO:0004930 | G-protein coupled receptor activity | 2.21E-03 |
| GO:0004871 | signal transducer activity | 5.38E-03 |

**Table 3.** Molecular function enrichment of gene expression changes in confluent HFFs upon tachyzoite infection.

| Category | Term | Padj |
| --- | --- | --- |
| Upregulated |  |  |
| GO:0003677 | DNA binding | 2.26E-13 |
| GO:0046982 | protein heterodimerization activity | 2.74E-11 |
| GO:0042393 | histone binding | 5.18E-11 |
| GO:0097159 | organic cyclic compound binding | 5.58E-08 |
| GO:1901363 | heterocyclic compound binding | 5.97E-08 |
| GO:0003676 | nucleic acid binding | 1.08E-05 |
| GO:0036094 | small molecule binding | 5.55E-05 |
| GO:0000166 | nucleotide binding | 1.07E-04 |
| GO:1901265 | nucleoside phosphate binding | 1.12E-04 |
| GO:0035575 | histone demethylase activity (H4-K20 specific) | 1.44E-04 |
| GO:0003688 | DNA replication origin binding | 4.10E-04 |
| GO:0003678 | DNA helicase activity | 5.19E-04 |
| GO:0005515 | protein binding | 9.61E-04 |
| GO:0097367 | carbohydrate derivative binding | 1.43E-03 |
| GO:0003697 | single-stranded DNA binding | 3.77E-03 |
| GO:0043168 | anion binding | 4.58E-03 |
| GO:0032451 | demethylase activity | 4.89E-03 |
| GO:0005488 | binding | 5.93E-03 |
| GO:0005200 | structural constituent of cytoskeleton | 9.56E-03 |
| GO:0032405 | MutLalpha complex binding | 1.03E-02 |
| GO:0050501 | hyaluronan synthase activity | 4.11E-02 |
| Downregulated |  |  |
| GO:0005539 | glycosaminoglycan binding | 3.81E-03 |
| GO:0005518 | collagen binding | 3.10E-02 |
| GO:0036122 | BMP binding | 3.10E-02 |
| GO:0048018 | receptor agonist activity | 3.32E-02 |

The majority of the host genes exhibiting altered expression 24 hours following *Toxoplasma* infection showed differences between proliferative and quiescent HFFs (Fig. 4B). We performed an Ingenuity Pathway Analysis (37) in order to determine which shared and distinct molecular and cellular functions were altered upon tachyzoite infection between confluent and subconfluent cells (Fig. 4C). We filtered our analyses to include all pathways that met a cutoff of Z score ≥ ± 2 in at least RFP or RNA datasets with Padj ≤ 0.05, and were not cancer-associated entries. Identified pathways were similarly represented for both the transcriptome and translatome levels independent of host cell confluency. Tachyzoite infection significantly repressed apoptotic and necrotic death pathways, whereas there was enhanced expression of genes involved in pathways associated with cell survival, viability, and migration. These findings are consistent with prior reports regarding the effects of tachyzoite infection on host cells (reviewed in (38, 39)). Of interest, an increased representation of transcription-related categories was observed specifically in the infected subconfluent cells, underscoring the level of transcriptional reprogramming that occurs in response to *Toxoplasma*. Infected confluent cells displayed an increase in cell cycle-associated pathways, suggesting that *Toxoplasma* infection induced these quiescent host cells to re-enter the cell cycle. Host cell cycle re-entry from quiescence has been reported by others and is characterized by an increased proportion of cells in S phase and a block in G2/M (8, 34, 40, 41). Taken together, these results demonstrate that quiescent and proliferating HFFs respond differently to tachyzoite infection. While both models of infection display gene expression changes consistent with reduced death and increased viability and motility upon infection, subconfluent cells predominantly show massive transcriptional rewiring and confluent cells are best characterized by re-entry into the cell cycle.

### Translational control in HFF cells infected with *Toxoplasma*

Recently, both the mTOR and ISR pathways were suggested to be modulated in the host upon *Toxoplasma* infection (9, 10). We used our Ribo-seq and RNA-seq data to examine the changes in mRNA translation that occur in quiescent and proliferating HFFs 24 hours post-infection. We compared the changes in steady-state mRNA levels to the changes in ribosome occupancy that occur in subconfluent and confluent HFFs harboring *Toxoplasma* (Fig. 5A-B). In subconfluent cells, 1,204 genes met a threshold of FDR = 0.01 whereas 632 genes passed the same filter in the confluent dataset (Table S3A-B). The genes in the subconfluent dataset clearly showed ribosome occupancy changes that tightly correlated (R^2^ = 0.9421) to the changes in steady state mRNA levels upon infection (Fig. 5A), suggesting that tachyzoite-induced changes to gene expression are driven in large part by changes in steady-state mRNA levels, likely via transcriptional control, in proliferating cells. In contrast, the correlation between transcript levels and ribosome occupancy in the confluent dataset was more widely dispersed (R^2^ = 0.8288), most noticeably due to a subset of genes in the upper left quadrant (Fig. 5B). Encoding mostly ribosomal proteins, these genes display modest decreases in mRNA levels while showing increased levels of translation, indicative of translational control acting in quiescent cells upon infection.

**Figure 5.**
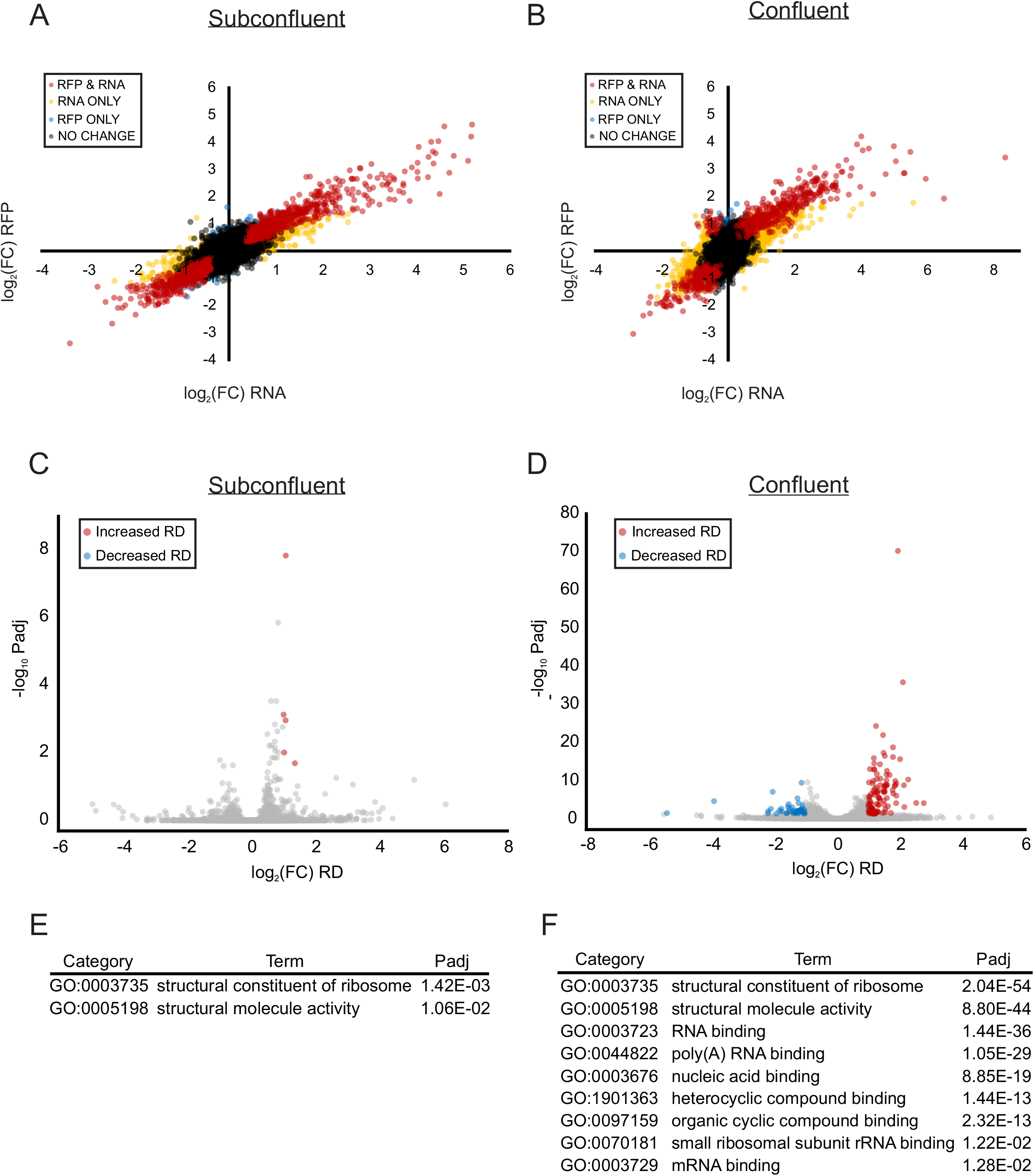
mTOR-coordinated translational control in HFFs 24 hours post-infection. A-B) Scatterplots of log_2_ fold changes of mRNA steady state (RNA) and ribosome footprint (RFP) libraries for (A) subconfluent and (B) confluent HFFs 24 hours post-infection. Colored genes are called by DESeq2 with an adjusted p value (Benjamini-Hochberg method) that meets a false discovery rate of 0.01. Red dots represent genes that change in mRNA and RFP abundance, yellow dots represent genes that change in mRNA abundance only, and blue dots represent genes that change in RFP abundance only. C-D) Scatterplot of log_2_ transformed fold change to ribosome density upon infection in (C) subconfluent and (D) confluent HFFs as a function of – log_10_ transformed Padj value. Significantly (FDR = 0.05) preferentially translated genes are in red while significantly translationally repressed genes are in blue. E-F) Gene ontology enrichment analysis of molecular functions associated with genes displaying increased ribosome density upon infection in subconfluent (C) and confluent (D) HFFs.

Although bulk translational capacity was largely unchanged in host cells upon infection (Fig. 1A), we analyzed the distribution of individual genes for changes in ribosome density. A total of 102 and 43 genes (FDR = 0.05) displayed increased or decreased ribosome density (log_2_FC ≥ ± 1), respectively, at our 24 hour post-infection time point in confluent HFFs (Fig. 5D, Table S4A). Gene ontology enrichment analysis (35) of the translationally repressed subset did not reveal any statistically significant (FDR = 0.05) enrichment terms associated with these genes. In contrast, the preferentially translated genes were strongly enriched for molecular functions relating to translation machinery including ribosomal proteins (Fig. 5F). These transcripts are canonically downstream targets for translational control acting through the mTOR complex 1 protein kinase (42). Increased activity of host mTOR complex 1 has been reported to occur in as little as two hours after *Toxoplasma* infection, leading to cell cycle progression, increased bulk translation, and the preferential translation of transcripts with 5’ oligopyrimidine tracts (7–9).

We also analyzed the dataset from subconfluent HFFs infected with *Toxoplasma* for 24 hours to determine which host genes displayed altered translational control (Table S4B). Applying a FDR of 0.05, only five genes were preferentially translated (log_2_FC ≥ 1), four of which were ribosomal proteins (Fig. 5C, E). In addition, although they did not meet our two-fold change cutoff, many other ribosomal proteins trended towards preferential translation in infected subconfluent HFFs (Table S4B). These results are consistent with a modest mTOR-like activation by *Toxoplasma*, which may be muted due to the strong basal mTOR activity in proliferating HFFs. There were no translationally repressed transcripts observed in the subconfluent dataset (Fig. 5C), suggesting that *Toxoplasma* does not induce a significant ISR in proliferating HFFs. Alternatively, *Toxoplasma* may induce an ISR earlier during infection in subconfluent HFFs that is resolved by the 24 hour time point. For example, in MEF cells, GCN2 was reported to be activated as early as two hours post-infection with *Toxoplasma* (10). The simultaneous Ribo-seq method reported here will serve as an invaluable tool to clarify host-parasite interactions in multiple cell types over the time course of infection.

## Conclusions

Our study demonstrates the feasibility of concurrently profiling host and parasite translation during infection using either proliferative or quiescent host cell models of infection. At 24 hours post-infection, we achieved nearly 25% representation of the *Toxoplasma* translatome, which was sufficient to allow identification of candidate uORFs for further interrogation. In addition, our results show that bulk translational capacity in tachyzoites remains unchanged in confluent or subconfluent host cells. We also demonstrated that both confluent and subconfluent HFFs exhibit gene expression patterns that are consistent with decreased cell death, increased survival, and cell cycle progression of previously quiescent cells. These gene expression patterns are consistent with prior observations of Toxoplasma-infected cells (8, 34, 38–41). Finally, our results revealed changes in translational control that are consistent with mTOR activation both in quiescent and, to a lesser extent, in proliferating HFFs. The ability to perform simultaneous ribosome profiling of intracellular pathogens within their host cells offers a new means to investigate host-parasite translational control in a wide-variety of other host cell types and culture conditions, further illuminating mechanisms underlying infection, drug activity, or development into latent bradyzoites in HFFs or other host cell backgrounds.

## Materials and Methods

### Host cell and parasite culture

Human foreskin fibroblasts (ATCC: SCRC-1041) were cultured in DMEM (Corning) supplemented with 10% fetal bovine serum (FBS; Atlanta Biologicals) without antibiotic/antimycotic. After reaching confluency, the monolayer was rinsed briefly with PBS, resuspended in trypsin, quenched with host cell media and passed into fresh flasks. Routine passage of *Toxoplasma gondii* strain RH (ATCC: 50174) was conducted as previously described (43). Briefly, confluent HFF monolayers were switched to DMEM media supplemented with 1% FBS without antibiotic/antimycotic and infected with parasites.

For all experiments using confluent host cells, HFF monolayers were allowed to reach confluency for one week prior to infection. Subconfluent HFFs were generated by subculturing host cells as described above 24 hours prior to infection. At this stage cells were 60% confluent. All experiments were conducted on host cells that were between passages seven and eleven. Infection of HFFs was conducted using freshly lysed parasites that were centrifuged at 3,400xg for 5 minutes and resuspended in fresh host cell media (e.g. with 10% FBS). Infection of confluent or subconfluent HFFs (MOI ~10) was conducted by adding prepared parasites to flasks without changing the media. Mock infection was conducted by adding the same volume of fresh media to uninfected flasks (200 μl of fresh media in 30 ml of pre-existing media). Parasites were incubated with host cells for 24 hours prior to sample collection for polysome or ribosome profiling.

### Polysome profiling

Polysome profiling was conducted as previously described (44). Briefly, cultured cells were incubated in 50 μg/ml cycloheximide (CHX) for 10 minutes, then washed in PBS containing CHX. The samples were lysed in cytoplasmic lysis buffer solution (20 mM Tris (pH 7.4), 100 mM NaCl, 10 mM MgCl_2_, 0.4% Nonidet P40, 50 μg/ml CHX), clarified by centrifugation at 12,000xg for 10 minutes and applied to a 10-50% sucrose gradient made in cytoplasmic lysis buffer without detergent. Gradients were subjected to centrifugation at 200,000xg for 2 hours at 4°C in a Beckman SW-41 Ti rotor. Polysome profiles were generated by applying the gradients to a Piston Gradient Fractionator (BioComp, Canada) and continuously reading the eluate at 254nm with an EconoUV monitor (BioRad, USA) paired with WinDaq software (DataQ instruments, USA). Polysomes were measured by calculating the area under the curve between the disome to the end of the polysomal region (outlined in Fig. 1D) and dividing by the total absorbance profiles.

### Ribosome profiling

Samples for ribosome profiling were generated as outlined in (12). Cytoplasmic extracts were generated as for polysome profiling with a few modifications: detergent was exchanged for 1% Triton X-100 and the buffer was supplemented with 25 units/ml of Turbo DNase I (Invitrogen). An aliquot of cytoplasmic lysate was immediately stored in TRIzol LS reagent (Ambion) for extraction of total RNA. The bulk of cytoplasmic lysate was incubated with 100 units of RNase I (Ambion) at 4°C for 1 hour while rotating. The amount of RNase I was empirically determined to optimize RFP generation by analyzing the polysome profiles of digested samples (Fig. 1D and data not shown). Sample digestion was quenched by the addition of 200 units of SUPERase·IN (Ambion). Digested samples were run on 10-50% sucrose gradients prepared as for polysome profiling with the addition of SUPERase·IN. The fractions corresponding to the monosome peak of centrifuged samples were collected using the same setup for polysome profiling paired with a Gilson fraction collector. RFPs and total RNA samples were collected from TRIzol. Total RNA was fragmented by alkaline hydrolysis. RFPs and fragmented RNA were collected by gel extraction from a 15% denaturing NuPAGE gel (Invitrogen).

The TruSeq Ribo Profile kit was used per manufacturer’s instructions to generate sequencing libraries for the confluent dataset. The subconfluent sequencing libraries were generated as previously described (12). Ribosomal RNA depletion was performed using the Ribo-Zero kit (Illumina) for all libraries with the modifications prescribed by (12). Single end 75bp reads were generated on a NextSeq system (Illumina).

### Sequencing data analysis

Annotated genomes and transcriptomes were downloaded from HostDB.org and ToxoDB.org (v37) (24). Libraries were depleted of reads aligning to human or *Toxoplasma* rRNA *in silico* using the bowtie algorithm (v0.1.0) (45) as implemented through the RiboGalaxy interface (46). Adapter sequences were removed from unaligned reads using the Clip function in the FASTX toolkit as implemented through the public Galaxy server (47).

Metagene plots were generated by aligning the libraries to the human or *Toxoplasma* transcriptomes with bowtie (45) and following the RiboSeqR pipeline (v1.0.5) (48) as implemented through the RiboGalaxy platform (46). All further analysis was conducted utilizing the publically assessable Galaxy server (47).

Differential gene expression, ribosomal density, and identification of translational controlled genes were conducted by first aligning the libraries to the human and *Toxoplasma* genomes with HISAT2 (v2.1.0) (49). Feature counts were obtained using the htseq-count algorithm (0.9.1) (50) using the -union option. Differential expression analysis was conducted with DESeq2 (v2.11.40.2) (26). Ribosomal density was determined by taking the quotient of RFPs and RNA-seq normalized read counts as obtained via DESeq2. Only genes that were detected at the transcriptome and translatome levels with a normalized read count of ≥ 5 were included for these analyses. Identification of translationally controlled gene expression determined by assessing changes in ribosome density between infected and mock-infected conditions using Riborex (51).

The ribosome density across each coding gene was calculated by determining the proportion of non-overlapping reads that segregated to 5’-UTR, CDS, or 3’-UTR feature annotations with the htseq-count algorithm (50) using the -intersection (strict) option. Only genes that had annotated 5’-UTRs, 3’-UTRs and CDS were included in the analysis. The total number of reads allocated to each feature was calculated with the formula (#5’reads / [#5’reads + #CDSreads + #3’reads]). Genes that harbored potential uORFs were identified by determining those that had two fold higher RFP reads in their annotated 5’-UTR compared to their CDS, with a minimum read count of at least 10 in both features.

### Gene and pathway enrichment analyses

Gene enrichment analysis of upregulated and downregulated genes as well as for preferentially translated or translationally repressed transcripts was performed with the goseq algorithm (v1.26.0) (35) as implemented through the publically assessable Galaxy server (47). Reported results were reduced in complexity using the Revigo webserver (52) and are limited to molecular functions that meet a FDR = 0.05.

Pathway enrichment analysis was performed using Ingenuity Pathway Analysis (37) (Qiagen). The analysis was restricted to those genes that met a normalized read count ≥ 5, a log_2_FC ≥ ± 1, and a Padj ≤ 0.01 as determined by DESeq2. Reported results are limited to the molecular and cellular functions that were not annotated as pertaining to cancer or tumors, and were disrupted by a Z score ± 2 with an associated Padj value ≤ 0.05.

### Accession number(s)

All datasets from this work have been made available in the NCBI GEO database (GSE129869).

## Acknowledgements

The authors would like to thank Kenny Carlson, Jocelyn Holmes, and Shun Liang for technical assistance as well as the Biology of Intracellular Pathogens Group at IUSM and other members of the Wek laboratory for helpful discussions.

This research was supported by a research grant from National Institutes of Health (AI124723 to W.J.S. and R.C.W.). The funders had no role in the study design, data collection and interpretation, or the decision to submit the work for publication.

